# Effects of pharmacological calcimimetics on colorectal cancer cells over-expressing the human calcium-sensing receptor

**DOI:** 10.1101/2020.07.09.195255

**Authors:** Luca Iamartino, Taha Elajnaf, Katharina Gall, Jacquelina David, Teresa Manhardt, Petra Heffeter, Michael Grusch, Sophia Derdak, Sabina Baumgartner-Parzer, Martin Schepelmann, Enikö Kallay

## Abstract

The calcium-sensing receptor (CaSR) is a ubiquitously expressed multifunctional G protein-coupled receptor. Several studies reported that the CaSR plays an anti-inflammatory and anti-tumorigenic role in the intestine, and that it is down-regulated during colorectal carcinogenesis. We hypothesized that intestine-specific positive allosteric CaSR modulators (type II calcimimetics) could be used for the treatment of intestinal pathologies. Therefore, the aim of this study was to determine the effect of pharmacological stimulation of CaSR on gene expression *in vitro* and on tumor growth *in vivo*.

We stably transduced two colon cancer cell lines (HT29 and Caco2) with lentiviral vectors containing either the *CaSR* fused to GFP or GFP only. Using RNA sequencing, RT-qPCR experiments and ELISA, we determined that CaSR over-expression itself had generally little effect on gene expression in these cells. However, treatment with 1μM of the calcimimetic NPS R-568 increased the expression of pro-inflammatory factors such as IL-23α and IL-8 and reduced the transcription of various differentiation markers in the cells over-expressing the CaSR. *In vivo*, neither the presence of the CaSR nor *p.o*. treatment of the animals with the calcimimetic cinacalcet affected tumor growth, tumor cell proliferation or tumor vascularization of murine HT29 xenografts.

In summary, CaSR stimulation in CaSR over-expressing cells enhanced the expression of inflammatory markers *in vitro*, but was not able to repress colorectal cancer tumorigenicity *in vivo*. These findings suggest potential pro-inflammatory effects of the CaSR and type II calcimimetics in the intestine.

## 1. Introduction

The calcium-sensing receptor (CaSR) is a G protein-coupled receptor (GPCR) belonging to the class C of the GPCR family. Its best studied function is to “sense” small variations in extracellular ionized calcium concentrations (Ca^2+^_ext_) in the bloodstream, maintaining it within a tight range *via* regulation of parathyroid hormone secretion in the parathyroid gland, bone resorption, and calcium excretion in the kidneys. The CaSR is the molecular target for cinacalcet (Mimpara/Sensipar), an FDA-approved positive allosteric modulator of the CaSR used to treat calcitropic disorders such as secondary hyperparathyroidism [1–3].

In addition to Ca^2+^, the CaSR is activated by other divalent and trivalent cations, cationic oligopeptides, polyamines and aminoglycoside antibiotics and is modulated by ionic strength, pH, L-amino acids and glutamyl peptides [3,4]. The CaSR has pleiotropic functions due to its propensity to bind various ligands, which can trigger different responses (ligand bias), depending on the specific cell context where the CaSR is expressed (system bias) [5,6]. The CaSR is expressed in many organs of the body, where it affects very different processes, such as neuronal differentiation [7], vessel tone and blood pressure [8], skin wound healing [9] insulin secretion [10,11] among many others [3]. In the gastrointestinal tract the CaSR regulates secretion of gastrointestinal hormones such as gastrin and cholecystokinin [12,13], controls fluid absorption [14], represses electrolyte secretion through the modulation of the enteric nervous system [15] and modulates intestinal barrier function and inflammation [16].

The CaSR is also involved in immune regulation where it can act both as a responder to inflammatory cytokines release and as a regulator of inflammation. Inflammatory cytokines upregulate CaSR expression in various cell types and animal models [17,18]. The CaSR regulates inflammation in a tissue-specific manner. In some tissues the CaSR promotes inflammation, *e.g.* in the respiratory tract, where it plays a pivotal role in asthma [19], and has pro-inflammatory role also in human adipose tissue [20]. In other tissues, such as in the intestine, the CaSR appears to suppress inflammation. The anti-inflammatory role of the CaSR in the intestine was first suggested by the studies on intestine-specific CaSR-knock-out mice that were more susceptible to dextran sulfate sodium (DSS)-induced colitis. Further studies showed that poly-L-lysine or glutamyl peptides, considered as orthosteric and allosteric CaSR ligands, ameliorated the symptoms of DSS-induced colitis [4,16,21,22]. However, none of these studies used the highly specific type II calcimimetics, although some used the calcilytic NPS 2143. Interestingly, in the only study that used cinacalcet for treating DSS-induced colitis, the anti-inflammatory effects could not be reproduced [23].

The multifaceted functions of the CaSR are reflected also in its dual role in tumorigenesis. In certain types of cancer such as in prostate and in breast cancer, the CaSR is up-regulated and promotes the formation of bone metastases [24,25]. In neuroblastoma and in colorectal cancer, however, it is down-regulated and it seems to function as a tumor suppressor [4,26–28]. Recent *in vitro* studies have reported that the CaSR counteracts colorectal cancer proliferation, inducing cell differentiation and apoptosis [26,29,30]. Consistent with this, global CaSR knock-out mice were more prone to develop pre-neoplastic lesions when challenged with azoxymethane (AOM)/DSS than their wild type counterparts [4,31].

As the CaSR has been suggested to act as a tumor suppressor in the colon, we hypothesized that targeting it may be beneficial for the treatment and prevention of colon cancer. Currently, there are no exhaustive studies of the effect of CaSR stimulation in CaSR expressing colon cancer cell lines or in *in vivo* models of colorectal tumors. Therefore, the aim of this study was to determine in an unbiased way, the effect of CaSR activation on gene expression in colon cancer cells and to test if restoration of CaSR expression affects tumor growth and if treatment with positive modulators of the CaSR strengthens this effect.

## 2. METHODS

### 2.1 Lentiviral transduction

We used two human colorectal cancer cell lines HT29 (ATCC, USA) and Caco2 (ATCC) in this study. 3×10^4^ HT29 cells and 3×10^5^ Caco2 cells were seeded in 6-wells plates and transduced with the Human Tagged ORF Clone lentiviral particles containing the pLenti-C-mGFP-P2A-Puro vector fused to the *CaSR* gene (RC211229L4V, OriGene, USA) (HT29^CaSR-GFP^ and Caco2^CaSR-GFP^). Cells transduced with the empty pLenti-C-mGFP-P2A-Puro vector (PS100093, OriGene), (HT29^GFP^ and Caco2^GFP^) were used as controls. HT29 cells were transduced with a multiplicity of infection (MOI) of 1 and Caco2 cells with a MOI of 10, in the presence of 8μg/ml of polybrene (TR-1003-G, Sigma-Aldrich, USA) to enhance transduction efficiency. The cells were incubated with the lentiviral particles for 20h, then left in normal culture medium for one day for recovery. Starting with the following day, the cells were selected with puromycin for 12 days at specific concentrations determined beforehand: 0.8μg/ml for HT29 and 10μg/ml for the Caco2 cells. The selection media were refreshed every other day and the cells were checked daily. The surviving cells were expanded and eventually frozen for long term storage.

### 2.2 Cell culture

Parental and transduced cells were cultured in Dulbecco’s Modified Eagle’s Medium (DMEM, GIBCO, USA) containing 10% fetal calf serum (FCS, GIBCO), 100U/ml Pen-Strep (GIBCO, USA), 2mM L-glutamine (GIBCO) and 10mM Hepes (GIBCO). Unless otherwise specified, calcium concentration was 2mM. All cells were periodically checked for mycoplasma contamination. Depending on the experiment, 70% confluent transduced cells were treated with 1μM of the positive CaSR modulator NPS R-568 (Tocris Bioscience, UK) or the negative CaSR modulator NPS 2143 (Tocris Bioscience) dissolved in dimethyl sulfoxide (DMSO). The maximum DMSO concentration in the culture medium never exceeded 0.1% (v/v). The transduced cells were periodically re-selected (once per month) with puromycin for 5 days.

### 2.3 FACS sorting and flow cytometry

The cells were detached with an enzyme-free dissociation buffer (1mM EDTA in Hanks’s balanced salt solution; HBSS) (GIBCO) at pH 7 and re-suspended in HBSS with 250μM HEPES and 1% bovine serum albumin (BSA) (Sigma-Aldrich, USA). The cells were sorted based on GFP intensity using “Moflo Astrios EQ” (Beckman Coulter, USA) FACS machine.

For measuring membrane expression of the CaSR *via* flow cytometry, the detached cells were re-suspended in phosphate-buffered saline (PBS) containing 0.1% BSA and polyclonal affinity purified anti-CaSR antibody, raised against the extracellular domain of the receptor (ImmunoGenes Kft., Hungary) and incubated for 40 minutes on ice. Alexa Fluor 647-conjugated anti-mouse IgG (A21235, ThermoFisher, USA) was used as secondary antibody for 30 minutes on ice in the dark. Fluorescence was detected by flow cytometry with FACSCanto (BD Biosciences, USA). Data were analyzed using the FlowJo V10 software (BD Biosciences).

### 2.4 RNA isolation and quantitative reverse transcription PCR (RT-qPCR)

The cells were harvested in TRIZOL (ThermoFisher) and the RNA was column-purified using the RNeasy Plus kit (QIAGEN, Germany) following the manufacturer’s protocol. The RNA was reverse transcribed using the High Capacity cDNA Reverse Transcription kit (Applied Biosystems, USA) following the manufacturer’s instructions. RT-qPCR was performed with Power SYBR Green PCR master mix (Applied Biosystems) using 7900HT Fast Real-Time PCR System (ThermoFisher). The amplicons were quantified with the relative quantification method using *ribosomal protein lateral stalk subunit P0 (RPLP0)* as reference gene. Primers against the specific targets are listed in the supplementary table S1. CaSR mRNA was quantified using the specific Taqman probe Hs01047795_m1 (ThermoFisher) using GoTaq Probe qPCR Master Mix (Promega, USA).

### 2.5 ELISA assay

The transduced cells were seeded in 6-well plates and upon reaching 70% confluency were treated either with 1μM NPS R-568, 1μM NPS 2143 or 0.1% DMSO (vehicle control) for 24h. The cell culture supernatant was collected and used for quantifying IL-8 secretion using the ELISA assay kit (# 88-8086-88, ThermoFisher). The assay was performed following the manufacturer’s protocol. Chemiluminescence was recorded with Infinite M200 Pro (TECAN, Austria) microplate reader.

### 2.6 Western Blot

Proteins were extracted using a lysis buffer containing 12mM Hepes, 300mM mannitol, 1mM EGTA, 1mM EDTA, 1%Triton X-100, and 0.1% SDS adjusted to pH 7.6, in the presence of 1mM n-ethylmaleimide, 1mM phenylmethanesulfonyl fluoride and protease inhibitor cocktail (P8340, Sigma-Aldrich). Protein extracts were clarified by centrifugation and then quantified using the Pierce BCA Protein Assay kit (ThermoFisher). 20μg of protein extracts were subjected to SDS-PAGE and then transferred onto a PVDF membrane. Immunoblotting was carried out using either anti-CaSR antibody (ImmunoGenes) or anti-Tubulin antibody (T5168, Sigma-Aldrich). HRP-conjugated anti-rabbit (111-035-003, Jackson ImmunoResearch, USA) and anti-mouse (115-035-003, Jackson ImmunoResearch) were used as secondary antibodies. Chemiluminescence was detected by a ChemiDoc MP Imaging System (Biorad, USA).

### 2.7 Cell immunofluorescence

Cells were cultured on 8-chamber slides (ThermoFisher) until 70% confluent, and then fixed in 3.6% formaldehyde. Endogenous fluorescence was quenched with 50mM NH_4_Cl, followed by blocking with 5% goat serum in PBS for 30 minutes. Cells were incubated overnight with anti-CaSR antibody (ImmunoGene) diluted in 0.1% BSA in PBS. As secondary antibody we used Alexa Fluor 647-conjugated anti-rabbit antibody (A21235, ThermoFisher). The nuclei were stained with 4′,6-diamidin-2-phenylindol (DAPI, Merck, Germany).

Confocal images were acquired using an inverted confocal microscope (Zeiss Invert Axio Observer.Z1) with a 63/1.4 oil objective. The fluorophores were excited at 405, 488 or 647nm, using a 405/488/647 multiline argon/krypton laser. Images were processed using ZEN 2.3 software (Zeiss, Germany).

### 2.8 RNA sequencing and data analysis

Sequencing libraries were prepared at the Genomics Core Facility, Medical University of Vienna, using the NEBNext Poly(A) mRNA Magnetic Isolation Module, the NEBNextUltra™ II Directional RNA Library Prep Kit for Illumina and the NEBNext® MultiplexOligos for Illumina® (Dual Index Primers Set 1) according to the manufacturer’s protocols (New England Biolabs, USA). Libraries were QC-checked on a Bioanalyzer 2100 (Agilent, USA) using a High Sensitivity DNA Kit for correct insert size and quantified using Qubit dsDNA HS Assay (Invitrogen). Pooled libraries were sequenced on a NextSeq500 instrument (Illumina, USA) in 1×75bp single-end sequencing mode. Approximately 28 million reads were generated per sample. Reads in fastq format were aligned to the human reference genome version GRCh38 [32] which was complemented with additional sequences for the lentiviral expression vector of CaSR, and with Gencode 29 annotations [33] using STAR aligner [34] version 2.6.1a in 2-pass mode. Reads per gene were counted by STAR, and differential gene expression was calculated using DESeq2 [35] version 1.20.0.

The changes in gene expression were calculated for pairwise group comparisons, and the statistical significance was calculated based on Wald test. False discovery rate adjusted p value (padj) was corrected using the Benjamini-Hochberg procedure.

Differentially expressed genes were shortlisted based on the adjusted *p* value (padj < 0.05) with a log_2_fold change either > 1 or < −1. The shortlisted genes were clustered and visualized in a heatmap using Cytoscape software version 3.7.1 [36]. The genes were clustered based on Pearson’s correlation using ClusterMaker 2.0 hierarchical cluster tool of Cytoscape [37]. Gene Ontology (GO) enrichment analysis was performed on the gene clusters using the online tool GOrilla (Gene Ontology enRIchment anaLysis and visuaLizAtion) [38].

### 2.9 Xenograft models

Eight weeks-old Balb/c severe combined immunodeficient (SCID) female mice (Envigo, Italy) were used for the animal experiments. The experiments were performed according to the regulations of the Ethics Committee for the Care and Use of Laboratory Animals at the Medical University of Vienna and the Austrian Ministry of Science (GZ: BMWFW-66.009/0006-WF/V/3b/2017).

Mice were separated in 4 groups of 8 mice each. Two groups (16 mice in total) were inoculated with 1×10^6^ HT29^CaSR-GFP^ cells and the other two groups with HT29^GFP^ cells suspended in 50μl serum-free medium. One group (8 mice) per cell line was treated *p.o*. either with cinacalcet, 10mg/kg bodyweight (Tocris Bioscience), the other with 20% of (2-hydroxypropyl)-β-cyclodextrin (Sigma-Aldrich), the vehicle in which cinacalcet was solubilized. The gavage started one week after the inoculation of the cells and was performed once per day on weekdays.

The mice were maintained in a pathogen-free environment and checked daily for distress; tumors were measured with a caliper and the tumor volume was calculated using the formula: tumor volume = length x width^2^. The mice were sacrificed when the first tumors ulcerated, which occurred 24 days post inoculation.

### 2.10 Tissue Immunohistochemistry and fluorescence quantification analysis

Tumors were fixed in 4% paraformaldehyde in PBS, dehydrated and embedded in paraffin and 4μm sections were cut. For immunofluorescence staining, the tumor sections were rehydrated and incubated in a steamer in 95 °C 10mM citrate buffer pH 6 for 20min for antigen retrieval. Tissues were permeabilized in 0.1–0.2% Tween 20 (Merck) and blocked in 5% goat serum (Abcam) containing 0.05% Tween 20 in PBS. For CD31 staining, the tissues were blocked in 1% BSA containing 0.1% Tween 20 in PBS. The sections were then incubated overnight at 4 °C with one of the following primary antibodies: rabbit polyclonal anti-mGFP (TA150122, OriGene, USA), mouse anti-hNuclei, clone 3E1.3 (MAB4383, Merck) and goat polyclonal anti CD31 (AF3628, R&D Systems). The sections stained for hNuclei were co-incubated with an eFluor 570-labelled rat monoclonal anti Ki-67 antibody (SolA15; ThermoFisher). After washing, the sections were incubated with either goat anti rabbit IgG AlexaFluor647 (A-21244, ThermoFisher), goat anti mouse IgG AlexaFluor647 (A-21235, ThermoFisher) or chicken anti-goat IgG AlexaFluor647 (A-21449, ThermoFisher) conjugated secondary antibodies. Sections were counterstained with DAPI (Merck) and mounted in Fluoromount G (Southern Biotech, USA). Images of the whole tumor sections were acquired using an automated microscope (Axioimager Z1, Zeiss, Germany) equipped with TissueFAXS hard- and software (TissueGnostics GmbH, Austria) using a 20x NeoFluar NA 0.5 objective (Zeiss). Image acquisition settings were kept constant for all tumor sections.

The acquired images were analyzed using the TissueQuest analysis software (TissueGnostics). Cells were detected based on segmentation of their DAPI stained nuclei. For detection of CD31, ring masks of a 2μm radius around the detected nuclei were used. For detection of hNuclei and Ki67, nuclear masks were used. Sections with fewer than 100,000 detected nuclei were not included in the analysis. Visible distortions and image artefacts were excluded from the analysis. The main tumor areas in the sections (*i.e.* excluding the surrounding stroma and skin) were segmented manually. Equal thresholds for positive/negative cells were set for all sections with the same staining and the number of positive and negative cells for each marker was determined. Where available, results from two sections of the same tumor were averaged as technical replicates. All analyses were performed by a scientist blinded to the genotypes / treatments of the tumors.

### 2.11 Statistical analysis

Data are presented as mean ± SD derived from 3 biological replicates for the *in vitro* tests and 8 mice per group for the *in vivo* experiments and the analysis of tumor biopsies. Statistical analysis was performed using GraphPad Prism version 7.0 (GraphPad, USA). The applied statistical tests are indicated in the captions of each figure.

## 3. Results

### 3.1 Generation and validation of the CaSR-overexpressing cell models

The expression of the CaSR is usually very low or undetectable in colorectal cancer cell lines, such as HT29 and Caco2. Therefore, we stably transduced HT29 and Caco2 cells with a lentiviral vector bearing either the *CaSR* fused with GFP (HT29^CaSR-GFP^ and Caco2^CaSR-GFP^), or an empty vector bearing only *GFP* (HT29^GFP^ and Caco2^GFP^).

Stably transduced cells were FACS-sorted *via* flow cytometry based on GFP intensity (fig. S1A-B). CaSR mRNA expression was verified by RT-qPCR (fig. 1A and S2A) and protein expression either with western blot (fig. 1B and S2B), or with immunofluorescence (fig. 1C and S2C). The GFP-staining in the HT29^CaSR-GFP^ and Caco2^CaSR-GFP^ cells was restricted primarily to the plasma membrane, whereas in the HT29^GFP^ and Caco2^GFP^ cells GFP fluorescence was evenly dispersed through the cytoplasm. CaSR staining with a CaSR-specific antibody co-localized at the plasma membrane with GFP (shown in yellow/orange in fig. 1C and S2C). We assessed CaSR recruitment to the plasma membrane with flow cytometry, and found a strong signal in HT29^CaSR-GFP^ and Caco2^CaSR-GFP^ cells compared with HT29^GFP^ and Caco2^GFP^ controls (fig. 1D and fig. S2D).

**Fig. 1.**
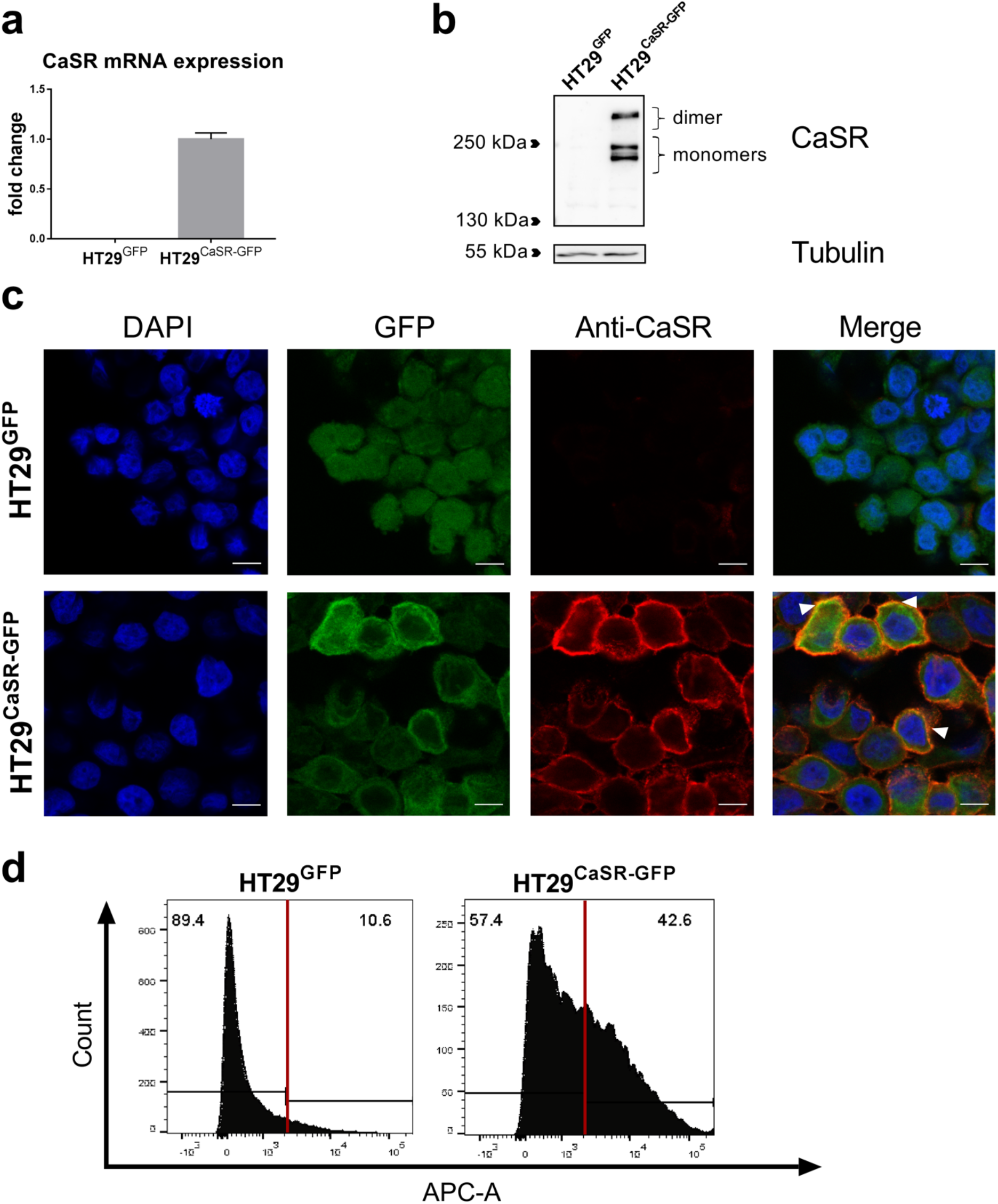
CaSR expression in transduced HT29 cells. CaSR mRNA quantification, values were normalized to the mean of CaSR expression in HT29^CaSR-GFP^ cells + SD (n=3) (**a**). Representative western blot for CaSR protein expression; in HT29^CaSR-GFP^ cells the CaSR monomers and the dimer corresponding to the CaSR-GFP fusion-protein were detected (**b**). Representative confocal images of GFP (green) and CaSR (red) (immuno-)fluorescence; white arrows indicate the points of co-localization (yellow/orange); scale bars represent 10μm (**c**). Representative histograms showing the percentages of CaSR-negative (left) and –positive (right) cells detected via flow cytometry, using the allophycocyanin (APC-A) channel (**d**).

We also tested CaSR functionality assessing the activation of the G protein subunit alpha q/11 (G_q/11_) – phospholipase C (PLC) – inositol 1,4,5-trisphosphate (IP3) cascade, which is coupled to the activated CaSR [3]. We measured inositol monophosphate (IP1) accumulation, a product of the degradation of IP3, under increasing concentrations of extracellular calcium. As expected, stimulation of HT29^CaSR-GFP^ cells with NPS R-568 caused a left shift of the concentration-response curve compared with the untreated control, whereas NPS 2143 induced a right shift, indicating that the transduced CaSR is functional and that its activation induces PI-4,5P_2_ breakdown to IP3 (fig. S3A). The EC_50_ for Ca^2+^_ext_ was 6.61mM ± 0.35, treatment with NPS R-568 lowered it to 5.30mM ± 0.7, while NPS 2143 increased it to 7.18mM ± 0.91.

In addition, we measured intracellular Ca^2+^ (Ca^2+^_int_) mobilization upon elevation of the extracellular calcium (Ca^2+^_ext_) concentration from 0.5 to 3.0mM, as a further test of functionality. Under these conditions, HT29^CaSR-GFP^ cells were responsive to Ca^2+^_ext_ whereas control cells HT29^GFP^ were not (fig. S3B). To further prove the functionality of the transduced CaSR, we tested the effects of CaSR-specific allosteric modulators. As shown in fig. S3C, Ca^2+^_int_ mobilization was increased by the CaSR positive modulator NPS R-568, and inhibited by the CaSR negative modulator NPS 2143.

### 3.2 The CaSR regulates gene expression

We compared whole transcriptome changes between HT29^CaSR-GFP^ and HT29^GFP^ cells treated for 48h with either vehicle (0.1% DMSO) or 1μM NPS R-568, to investigate whether CaSR over-expression and/or CaSR stimulation affect transcription. Differentially expressed genes were plotted in a heat-map (fig. 2A). Already over-expressing the CaSR changed the expression of several genes compared with the HT29^GFP^ controls. This effect was further enhanced by the treatment of the HT29^CaSR-GFP^ cells with NPS R-568, revealing a profound effect of CaSR stimulation on the transcriptome. The treatment with NPS R-568 caused significant changes in gene expression only in HT29^CaSR-GFP^ cells, but not in HT29^GFP^ cells, demonstrating that the effects of the drug were mediated *via* the CaSR (fig. 2A).

**Fig. 2.**
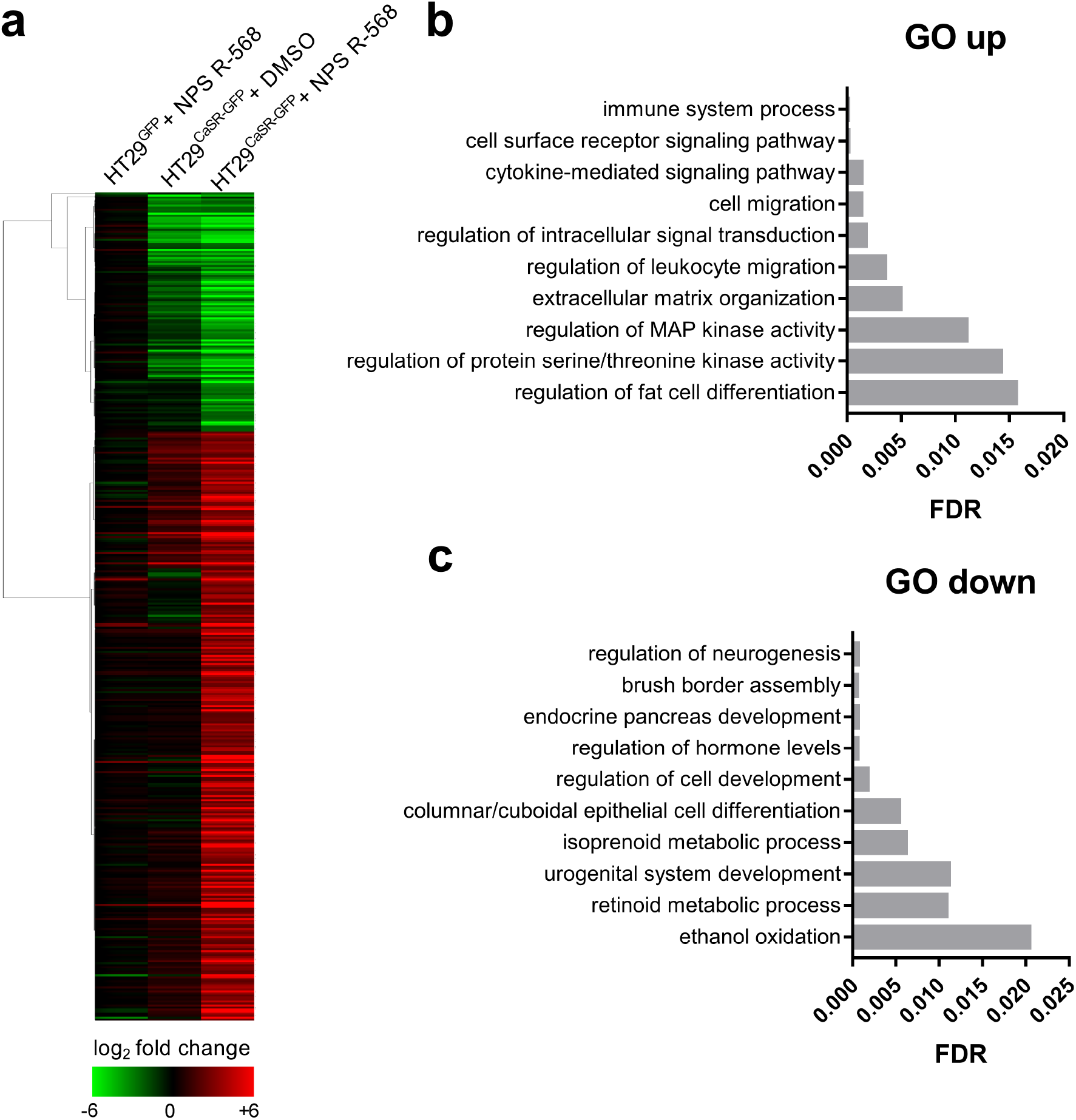
RNA-seq analysis. The heat-map shows the differentially expressed genes (the up-regulated in red and the down-regulated in green) after treatment of HT29^GFP^ and HT29^CaSR-GFP^ cells for 48 hours with NPS R-568 or with DMSO. The genes were clustered based on their expression values using Pearson’s correlation and compared with the expression of vehicle-(DMSO) treated HT29^GFP^ cells using a green-red gradient scale. The expression values plotted in the heat-map represent the means of three biological replicates (**a**). Panels **b** and **c** show the Gene Ontology (GO) enrichment analysis performed on the up-regulated and on the down-regulated gene subsets of the HT29^CaSR-GFP^ + NPS R-568 group. The enriched pathways are plotted against their FDR q-val (adjusted p value for False Discovery Rate).

GO enrichment analysis of differentially expressed genes indicated that CaSR stimulation with NPS R-568 up-regulated genes belonging to pathways related to immune responses, “extracellular matrix organization” and membrane receptor signaling including “regulation of mitogen-activated protein (MAP) kinase activity” and “regulation of protein serine/threonine kinase activity” (fig. 2B). Furthermore NPS R-568 inhibited the expression of differentiation factors, *i.e.* those linked to the “columnar/cuboidal epithelial cell differentiation” pathway (fig. 2C), or those involved in the “brush border assembly” complex which is essential for the maturation of microvilli in intestinal cells [39].

### 3.3 CaSR mediated up-regulation of inflammatory markers

To validate the RNA-seq results we assessed the mRNA expression of inflammation-related factors after treating the cells with 1μM of the positive and negative CaSR modulators for 24 and 48h. Interleukin-1 alpha (IL-1α) mRNA was up-regulated in HT29^CaSR-GFP^ cells treated with NPS R-568 (2.3-fold at 24h and 3.4-fold at 48h) compared with the DMSO control (fig. 3A). Similarly, NPS R-568 up-regulated IL-8 mRNA level 2-fold after 24h and 2.6-fold after 48h treatment (fig. 3B), and also increased significantly the expression of other cytokines, *e.g.* IL-23α, C-C motif chemokine ligand 20 (CCL20), colony stimulating factor 1 (CSF1) and interleukin-34 (IL-34), (fig. 3C-F). We observed similar effects in Caco2^CaSR-GFP^ cells, where NPS R-568 treatment promoted the expression of the same pro-inflammatory cytokines, with the exception of IL-34 (fig. S4A-F).

**Fig. 3.**
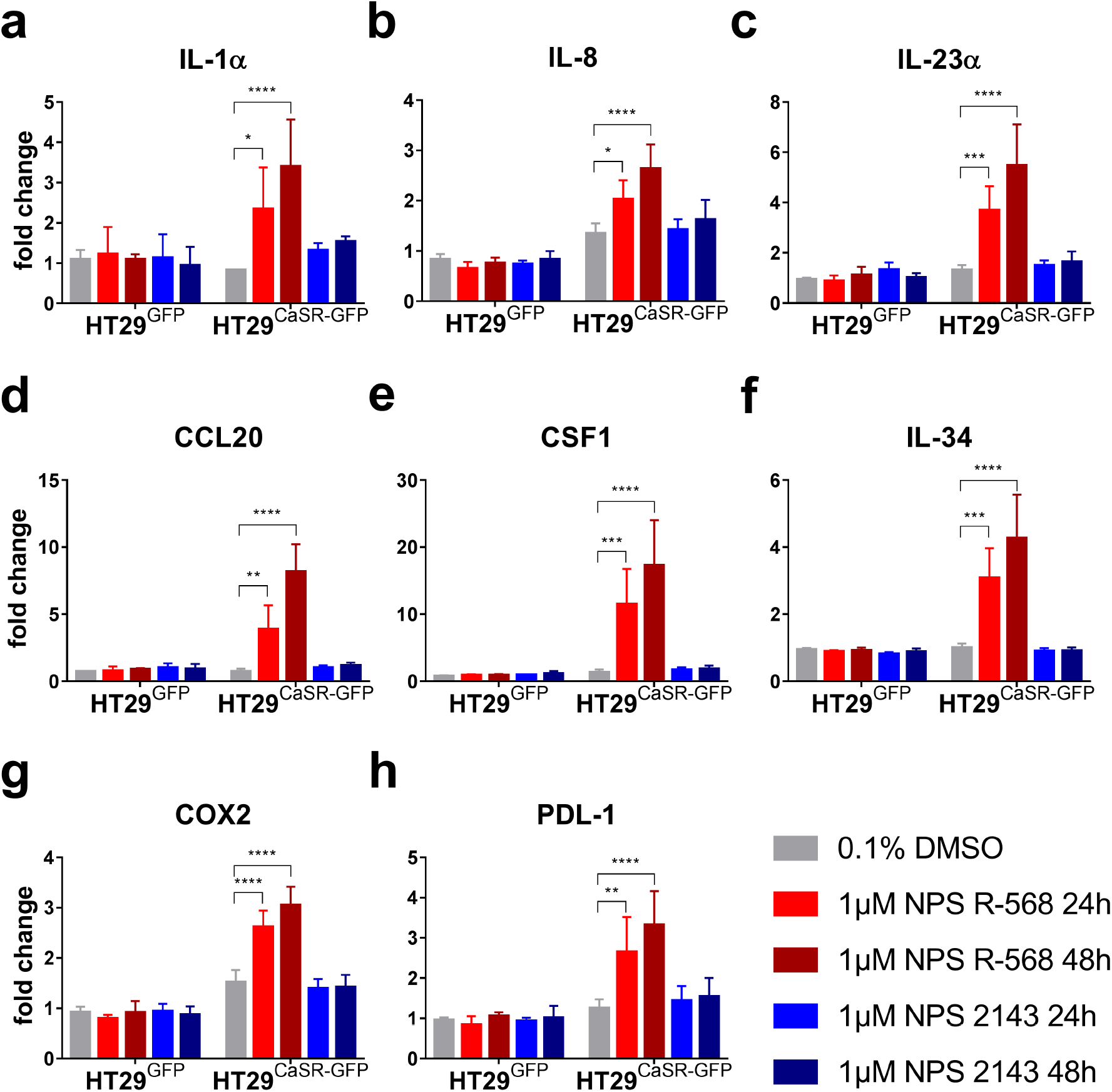
Expression of inflammatory markers in transduced HT29 cells. RT-qPCR analysis of the mRNA level of IL-1α (**a**), IL-8 (**b**), IL-23α (**c**), CCL20 (**d**), CSF1 (**e**), IL-34 (**f**), COX2 (**g**) and PDL-1 (**h**). All values were normalized to the untreated HT29^GFP^ cells as control. Data are presented as mean + SD (n=3). Statistical analysis was performed using two-way ANOVA with Dunnett’s post-hoc test, **** p<0.0001; *** p<0.001; ** p<0.01; * p<0.05.

We also assessed the mRNA level of inflammation-related factors that play an important role in tumorigenesis. Cyclooxygenase 2 (COX2) is frequently up-regulated in colorectal cancer [40]. In HT29^CaSR-GFP^ cells COX2 mRNA level was higher than in HT29^GFP^ cells and increased further (around 2-fold) in response to NPS R-568 treatment for 24h and 48h (fig. 3G). Similarly, in Caco2^CaSR-GFP^ cells COX2 mRNA level increased significantly after treatment with NPS R-568 (fig. S4G). Programmed cell death 1 ligand 1 (PDL-1) is a well-studied factor involved in immune check-point resistance, commonly over-expressed in malignant tumors that escape immune surveillance [41]. NPS R-568 increased the expression of PDL-1 in HT29^CaSR-GFP^ cells 2-fold after 24 h and 3-fold after 48h incubation (fig. 3H). NPS R-568 increased PDL-1 mRNA also in Caco2^CaSR-GFP^ cells but only after 48h (fig. S4H).

In order to confirm that NPS R-568 induces expression of the inflammatory markers in CaSR over-expressing cells not only at the mRNA but also at the protein level, we assessed secretion of the pro-inflammatory cytokine IL-8 by ELISA. Stimulating the CaSR with NPS R-568 for 24h in HT29^CaSR-GFP^ and in Caco2^CaSR-GFP^ cells enhanced IL-8 secretion 3.5-fold (fig. 4A) and 8-fold (fig. 4B) respectively, compared with the untreated controls.

**Fig. 4.**
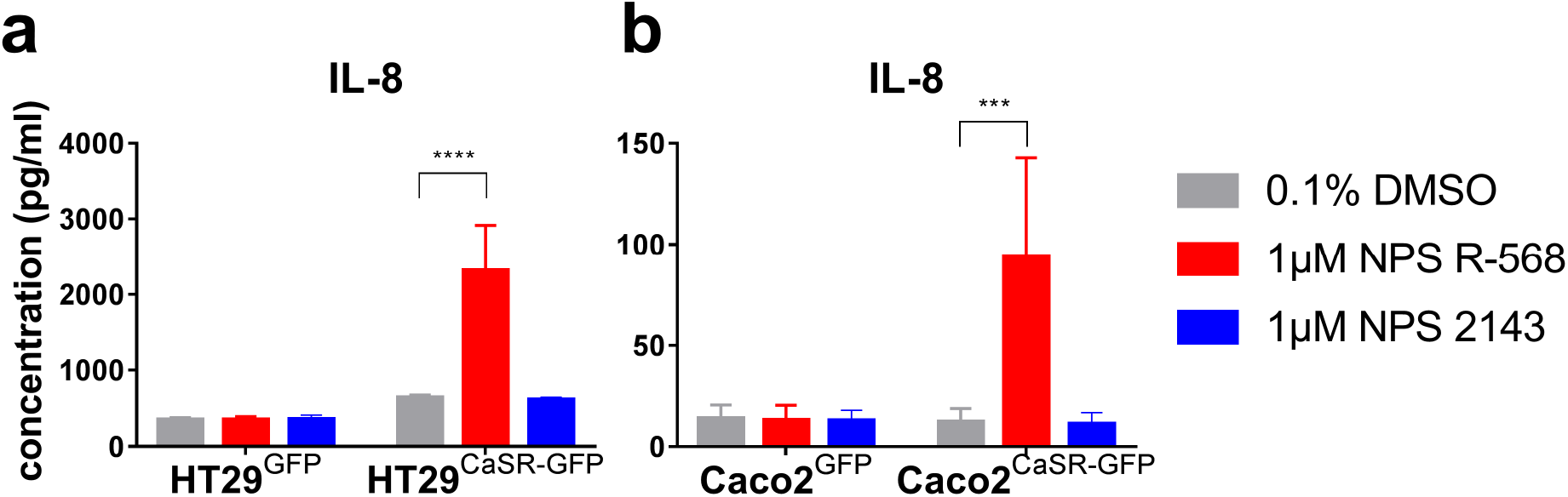
IL-8 secretion in transduced cells. ELISA assay for IL-8 secretion in the supernatant of the transduced HT29 (**a**) and Caco2 (**b**) cells after 24h treatment. Data are presented as mean + SD (n=3). Statistical analysis was performed using two-way ANOVA with Dunnett’s post-hoc test, **** p<0.0001; *** p<0.001; ** p<0.01; * p<0.05.

The negative modulator NPS 2143 did not affect the expression of any of the inflammatory factors tested when compared with the DMSO control in either of the cell lines tested.

### 3.4 CaSR influenced the transcription of intestinal markers

The RNA-seq results suggested that several intestinal differentiation markers were down-regulated due to the overexpression of the CaSR or by NPS R-568 treatment. To validate the data, we measured mRNA expression after 24 and 48h treatment with either 1μM of NPS R-568 or 1μM of NPS 2143. We measured the expression of several members of the brush border assembly complex and found that myosin 7 B (MYO7B), Ankyrin repeat and sterile alpha motif domain containing 4B (ANKS4B) and Usher syndrome 1 c (USH1C) were down-regulated in HT29^CaSR-GFP^ cells when compared with control HT29^GFP^ cells (fig. 5A-C). Similarly, HT29^CaSR-GFP^ cells expressed significantly lower levels of caudal type homeobox 2 (CDX2), a key factor for enterocyte differentiation [42] (fig. 5D); neurogenin3 (NEUROG3), which is essential for enteroendocrine cell differentiation [43] (fig. 5E); atonal BHLH transcription factor 1 (ATOH1); delta-like canonical NOTCH ligand 1 (DLL1) and 4 (DLL4), which are factors that control NOTCH signaling and regulate secretory cell maturation and maintenance [44] (fig. 5F-H), than the HT29^GFP^ cells. In the HT29^CaSR-GFP^ cells NPS R-568 further reduced the expression of these genes, but only moderately. The negative allosteric modulator NPS 2143 did not change the expression of any of the differentiation factors tested.

**Fig. 5.**
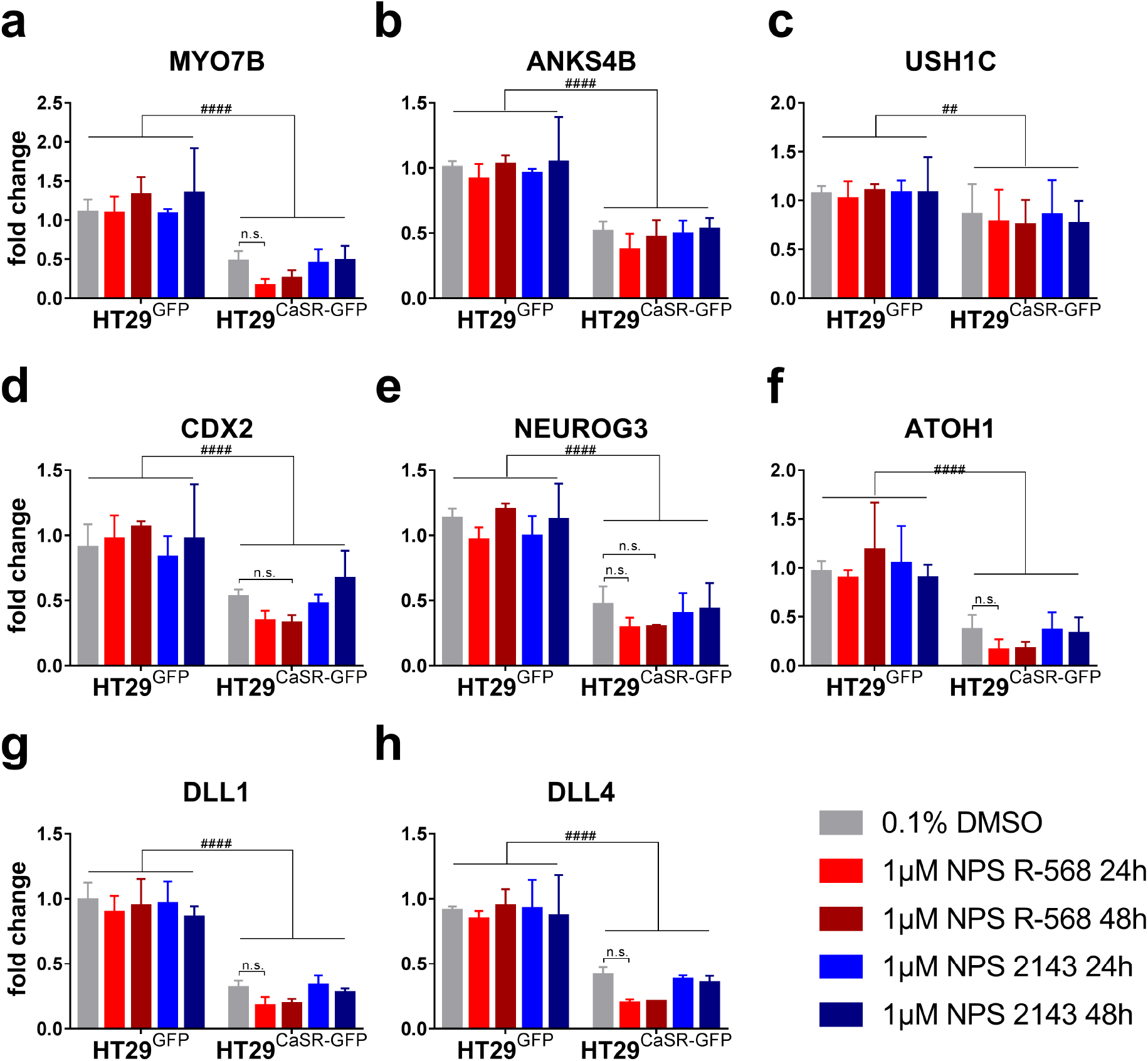
Expression of intestinal markers in transduced HT29. RT-qPCR analysis of the mRNA level of MYO7B (**a**), ANKS4B (**b**), USH1C (**c**), CDX2 (**d**), NEUROG3 (**e**), ATOH1 (**f**), DLL1 (**g**) and DLL4 (**h**). All values were normalized to the untreated HT29^GFP^ cells. Data are presented as mean + SD (n=3). Statistical analysis was performed using two-way Anova with Dunnett’s post-hoc test, ^####^ p<0.0001; ^###^ p<0.001; ^##^ p<0.01; ^#^ p<0.05; n.s.= not significant.

Whereas in transduced HT29 cells the expression of intestinal markers was inhibited primarily in response to over-expression of the CaSR with only minor additional effects in response to NPS R-568, in Caco2 cells the expressions of very few of the markers tested were affected by CaSR over-expression (Fig. S5). One exception was ATOH1, whose expression was significantly higher in Caco2^CaSR-GFP^ cells than in Caco2^GFP^ cells (fig. S5F), albeit the expression level was extremely low in both. The only other significant changes in the expression of intestinal markers in Caco2^CaSR-GFP^ cells were due to the treatment with NPS R-568, which reduced the expression of CDX2 (fig. S5D), DLL1 (fig. S5G) and DLL4 (fig. S5H). With respect to the brush border assembly complex, NPS R-568 had apparently contradictory effects, up-regulating MYO7B (fig. S5A) and down-regulating USH1C (fig. S5C) when compared with untreated controls.

### 3.5 Impact of CaSR expression and activation on tumor growth in a mouse xenograft model

We next tested whether over-expressing the CaSR and activating it with the positive CaSR modulator cinacalcet can counteract tumor growth *in vivo*. Therefore, we inoculated either HT29^CaSR-GFP^ cells or HT29^GFP^ cells under the skin of SCID Balb/c mice to generate sub-cutaneous tumor xenografts. One week after the inoculation, we started treating the animals with either cinacalcet (10mg/kg body weight) or vehicle (20% cyclodextrin) *via* gavage once per day. Three weeks after inoculation of the xenografts we were obliged to stop the experiment because the tumors in some mice had started to ulcerate. Ulcerations appeared to occur randomly, being independent of CaSR expression level or treatment. Tumor growth rates were comparable among all tested groups (fig. 6A) and tumor weights on the day of sacrifice were also similar (fig. 6B).

**Fig. 6.**
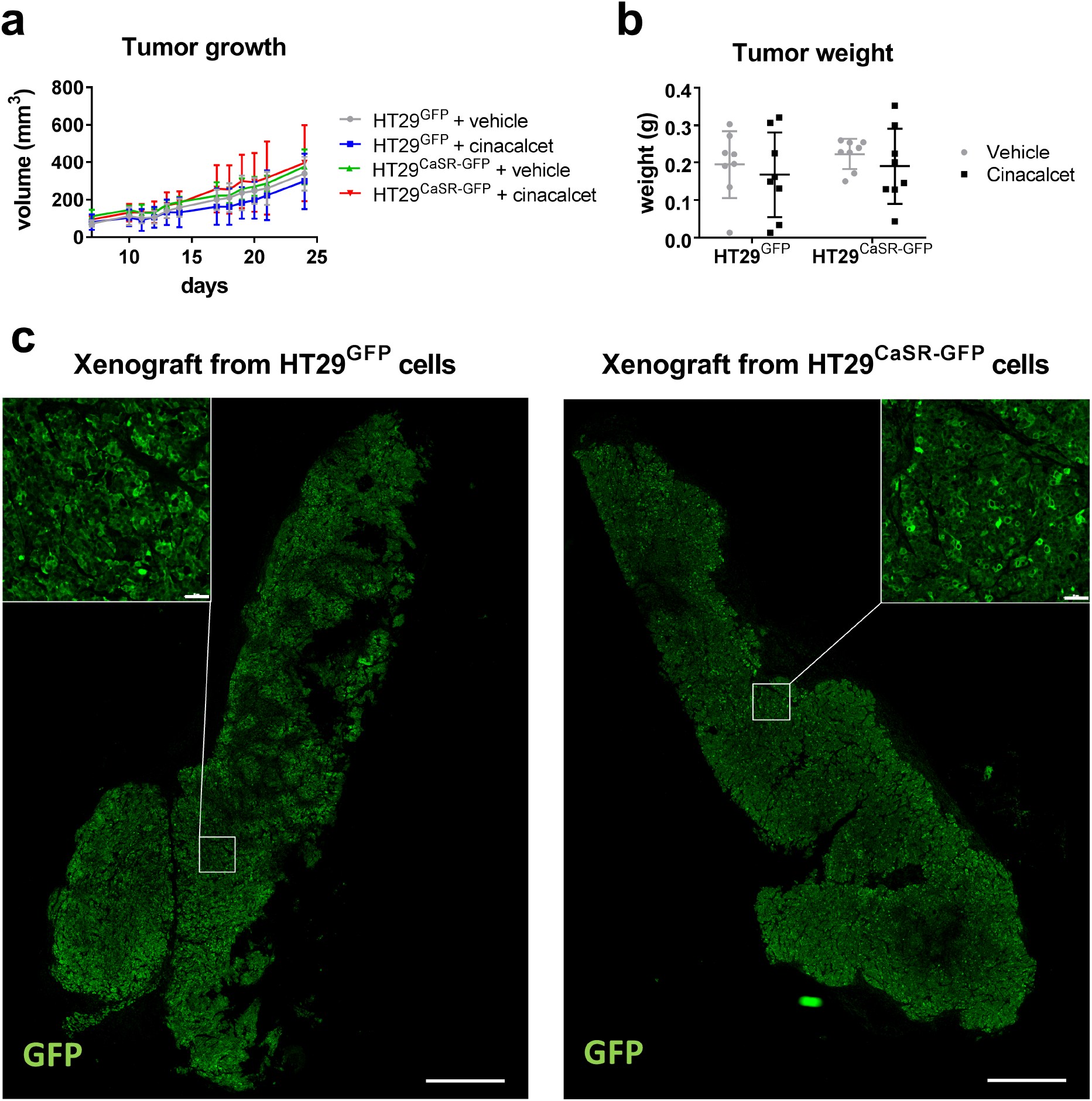
Tumor xenografts derived from the transduced HT29. Effect of CaSR expression and cinacalcet treatment on tumor volume, calculated using the formula (length × width^2^)/2 (**a**) and on tumor weight (**b**). (**c**) Representative images showing immunofluorescence staining for GFP (green) in tumors derived from HT29^GFP^ (left panel) and from HT29^CaSR-GFP^ (right panel) cells. Nuclei were counterstained with DAPI (blue). Each group had n=16 animals. Error bars represent standard deviations. Scale bars of the whole tumor figure represent 1 mm, while the scale bars within the insets represent 50μm.

The tumor cells retained the transduced GFP-linked constructs, as determined *via* immunofluorescence on tumor sections (fig. 6C). Neither over-expression of the CaSR, nor its activation with NPS R-568 affected tumor angiogenesis, as shown by staining with a specific antibody against murine platelet and endothelial cell adhesion molecule 1 (PECAM1 / CD31) (fig. 7A). Quantitative image analysis showed that the percentage of CD31-positive murine cells was similar in the tumors from the different groups (fig. 7C). We also assessed the expression of the proliferation marker Ki67, which was co-stained with a marker for human nuclei, in order to distinguish proliferating human tumor cells from infiltrating mouse cells (fig. 7B). The expression intensity of Ki67 in human cells and the percentage of proliferating human cells in all tumors was similar (fig. 7D), suggesting no effect of the CaSR level or treatment on the proliferative potential of the tumors. These results were supported by *in vitro* data, where neither over-expression of the CaSR nor its modulation with the allosteric modulators affected cell proliferation (fig. S6A and C) or viability (fig. S6B and D).

**Fig. 7.**
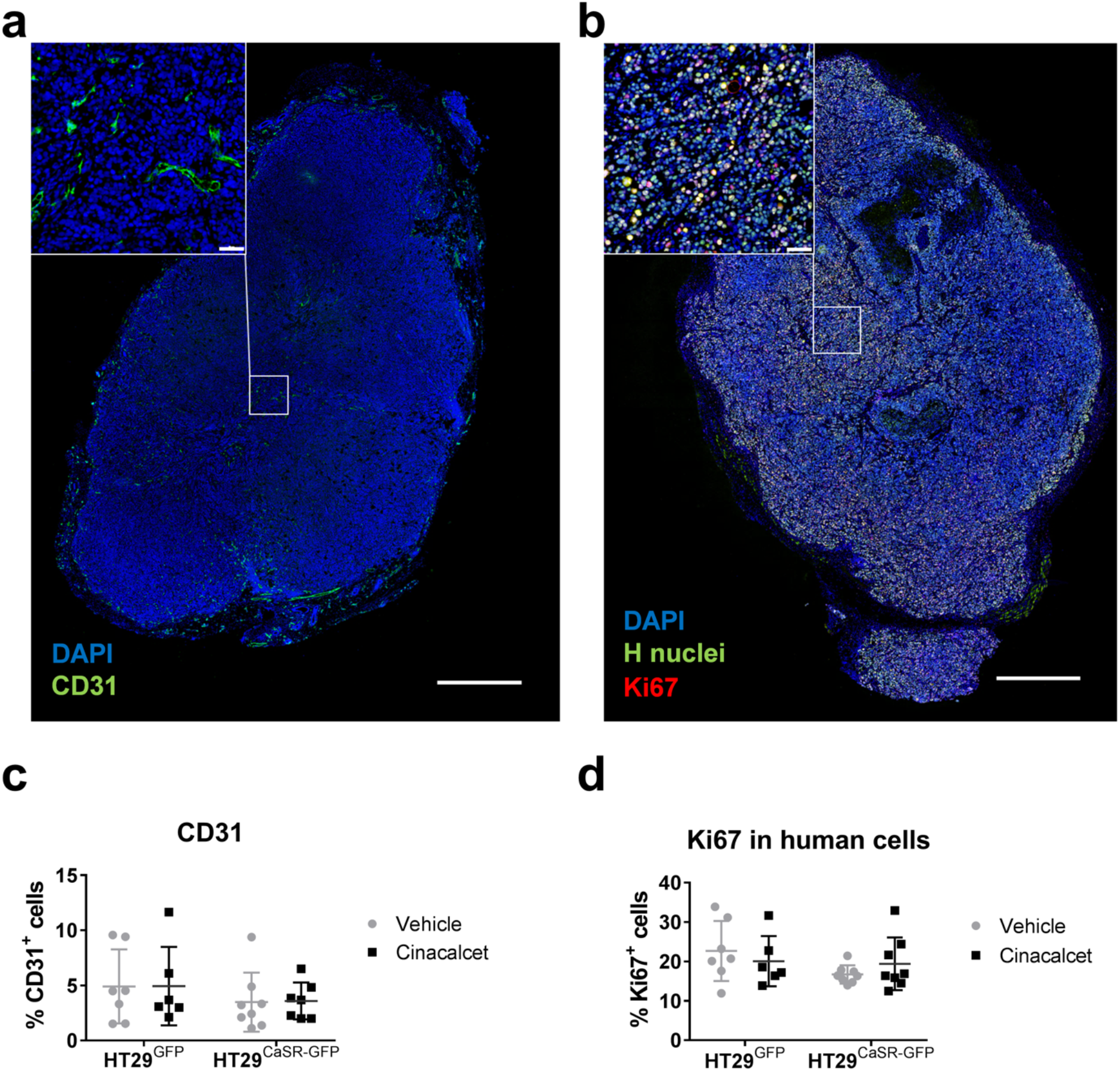
Angiogenesis and cell proliferation of tumor xenografts. Representative images showing immunofluorescence staining of the angiogenesis marker CD31 (green) (**a**) human nuclei (green) and the proliferation marker Ki67 (red) (**b**) in the tumors. Mouse and human nuclei were counterstained with DAPI (blue). Percentage of CD31 positive cells inside the tumors, the surrounding tumor stroma was excluded manually (**c**); percentage of Ki67-positive human cells (**d**). Each group had n=8 animals. Tumor sections with fewer than 100,000 cells were excluded from the quantitative image analysis. Error bars represent standard deviations. Scale bars of the whole tumor figure represent 1 mm, while the scale bars within the insets represent 50μm.

## Discussion

In this study we provide evidence that in colon cancer cells, which stably over-express the CaSR, activating the CaSR has a pro-inflammatory effect. We also show for the first time that neither over-expression of the CaSR, nor its pharmacological activation with the highly specific calcimimetic cinacalcet were able to inhibit or reduce tumor growth in a xenograft tumor model.

In certain tissues the CaSR promotes inflammation, such as in leukocytes [45–47], adipose tissue [48] and in the respiratory tract [19,49]. Moreover, pharmacological inhibition of the CaSR has been proposed as new therapeutic approach for the treatment of airway hyperresponsiveness and thus of asthma [19]. In the intestines however, the role of the CaSR is not completely clear, as several other studies suggested that there the receptor has anti-inflammatory effects, while our data suggest the opposite.

The Gene Ontology analysis of our cell model data set revealed that the genes up-regulated in HT29^CaSR-GFP^ cells after NPS R-568 treatment were significantly enriched in pathways involved in the inflammatory response. As this increase in the expression of pro-inflammatory markers was not seen in HT29^GFP^ cells, the CaSR appears to have mediated this inflammatory effect. This implies that activation of the CaSR could promote inflammation also in the intestine. In both HT29^CaSR-GFP^ and Caco2^CaSR-GFP^ cells stimulating the CaSR with the calcimimetic NPS R-568 increased the expression of several pro-inflammatory cytokines such as IL-1α, IL-8, IL-23α, CCL20 and CSF1, which are commonly up-regulated in IBD [50–52]. This suggests that the CaSR is involved in the pathogenesis of IBD. Indeed, this is in line with our recently published study which found that in a DSS-induced colitis model cinacalcet enhanced systemic inflammation and that inhibiting the CaSR with NPS 2143 ameliorated colitis symptoms and reduced immune cell infiltration in the mucosa [23].

The CaSR can interact directly with the immune system by activating the inflammatory response in monocytes through the NLR Family Pyrin Domain Containing 3 (NLRP3) inflammasome [46]. In monocytes, extracellular calcium activated NLRP3 and lead to increased levels of inflammatory cytokines like IL-1β and its target IL-6 [53,54]. However, in our RNA-seq dataset the expression level of NLRP3 was relatively low and was not significantly affected either by the overexpression of the CaSR or the treatment with NPS R-568 (data not shown) [55]. Furthermore, both IL-1β and IL-6 transcripts were barely detectable, suggesting that the activation of the CaSR in these cancer cells does not affect the inflammasome.

NPS R-568 treatment stimulated also the expression of pro-inflammatory factors which are well-known oncogenic markers, such as COX2 and PDL-1. COX2 is over-expressed in about 85% of colorectal adenocarcinomas and it is responsible for the synthesis of prostaglandins that play a major role during the inflammatory response, hence in colitis and in colitis-associated colorectal cancer [40,56]. In our model, the CaSR stimulated COX2 expression, as it does also in kidney cells [57] and in inflamed lung [49]. PDL-1 is an immune checkpoint protein and it is over-expressed by cancer cells to escape immune surveillance, inhibiting lymphocyte activation [41]. We found that CaSR stimulation with NPS R-568 enhanced the expression of PDL-1, potentially acting to suppress an effective anti-cancer immune response. Interestingly, a recent meta-analysis demonstrated an association between the use of immune check point inhibitors and the development of hypocalcemia [58], while several case reports have shown that inhibiting PDL-1-mediated immune tolerance-caused hypocalcemia and hypoparathyroidism, two key clinical features of CaSR hyperactivity [59,60]. These studies further substantiate a possible link between the CaSR and PDL-1, although the mechanisms underlying these effects are unclear and require further study.

GO analysis of our RNA-seq dataset revealed that, in addition to the upregulation of inflammatory processes, the CaSR inhibited molecular processes related to intestinal differentiation as well as pancreatic and neuronal development, reducing the expression of several transcription factors involved in these processes. Despite the different cellular contexts (intestinal *vs* pancreatic *vs* neuronal), key regulators of these mechanisms are shared. For example, the transcription factors enriched in the GO analysis such as NEUROG3, ATOH1, DLL1 and DLL4 control both neurogenesis and intestinal development [43,44,61,62]. Moreover, NEUROG3 together with insulinoma-associated 1 (INSM1) and paired box 4 (PAX4), which were also enriched in the GO analysis, are required for the development of both enteroendocrine cells and pancreas β-cells [63–66], although INSM1 is also a marker of neuroendocrine tumors [67,68]. Our results suggesting that the CaSR inhibits pathways involved in endocrine pancreas development, might explain why mice with CaSR activating mutations have reduced pancreatic islet cell mass and suffer from glucose intolerance [11].

We found that the CaSR had opposing effects on the expression of certain intestinal differentiation markers in the two analyzed cell lines. In HT29^CaSR-GFP^ cells, over-expressing the CaSR suppressed differentiation towards the epithelial and secretory lineage, *i.e.* it suppressed the expression of CDX2, which is important for the maturation of absorptive enterocyte cell [42], and the expression of NEUROG3 and ATOH1, which promote the differentiation of secretory cells [43,44]. However, over-expressing the CaSR inhibited also the expression of two main Notch pathway ligands, DLL1 and DLL4 that are inhibitors of secretory differentiation and support intestinal stem cell homeostasis [44]. Moreover we found no significant changes in the expression of secretory goblet cell markers such as mucin 2 (MUC2), or trefoil factor peptides [69] in the RNA-seq dataset (data not shown), thus it was not clear whether the CaSR promoted or inhibited the commitment of secretory cells lineage. In HT29^CaSR-GFP^ cells, CaSR over-expression suppressed also the expression of the brush border assembly complex, which promotes the formation and organization of microvilli [39,70], while the inhibitory effect of NPS R-568 was not statistically significant. In contrast, in Caco2^CaSR-GFP^ cells, the overexpression of the CaSR had no clear effect on the expression of these factors, but the treatment with NPS R-568 inhibited the transcription of the intestinal differentiation markers CDX2, DLL1 and DLL4. Both HT29 and Caco2 cell lines derive from colorectal adenocarcinomas, however they are very different in terms of genetic background and differentiation level [71,72]. Our data suggest that the cells genetic background and their pre-existing differentiation state affected the nature of the CaSR response.

Taken together, in the two different cell lines, overexpression and activation of the CaSR had similar effects on inflammatory markers, but appeared to have contradictory effects on cell differentiation, indicating that these effects are unlikely to be important in intestinal physiology and more likely affect pathophysiology.

Neither the presence of the CaSR nor its stimulation with cinacalcet affected growth, proliferation or angiogenesis of HT29^CaSR-GFP^-derived tumor xenografts in mice.

The results of our study are in contrast with previous publications, which have suggested a protective role for the CaSR in the intestine (as reviewed in [4]). CaSR knock-out mouse models indicated that the CaSR prevents the development of intestinal malignancies [16,31]. However, it is possible that the CaSR preserves epithelial integrity or regulates the tumor microenvironment, rather than acting directly on malignant cells. Moreover, CaSR knock-out mice may develop compensatory mechanisms that interfere with normal cell homeostasis and thus induce enhanced sensitivity to carcinogenic and pro-inflammatory insults [16,31].

Some *in vitro* studies using colorectal cancer cell lines reported that the CaSR inhibited the expression of IL-8 and IL-6, and induced the expression of the anti-inflammatory cytokine IL-10 [21,22]. An explanation for these discrepancies with our data could be that in these reports the CaSR was studied in cell lines in which endogenous expression was limited [21,22,73–76]. In the current study, however, we over-expressed the CaSR with an exogenous construct. The cellular model we created expressed functional CaSR, as revealed by intracellular calcium mobilization assay and by IP1 accumulation assay. Moreover, the transfected CaSR was efficiently transported and localized to the cell membrane. In addition, different transfection/transduction systems might have led to different results depending on the level of expression or subcellular localization of the receptor [26,77,78].

Over-expression of the CaSR constitutes a limitation of the experimental system as high levels of CaSR mRNA and protein may interfere with normal cell homeostasis. Finding a way to re-establish endogenous expression of the CaSR in colorectal cancer cells or to increase the level of CaSR expression in pre-cancerous cells would more accurately clarify the role of the CaSR in intestinal inflammation and tumorigenesis. Moreover, to elucidate whether the CaSR and its positive modulators induce intestinal inflammation, studies using biopsies from healthy and inflamed intestinal tissue or colonic organoids are required. This would help clarify whether the CaSR might be a new pharmacological target for the treatment of inflammatory bowel diseases.

Even though it is unclear whether these observations are connected, clinically used type II calcimimetics such as cinacalcet or etelcalcetide are known for causing gastrointestinal side effects such as nausea and vomiting [79]. Furthermore, cinacalcet can cause gastrointestinal bleeding, mostly in the upper digestive tract [80]. Our results together with this evidence from pharmacovigilance suggest that stimulating the CaSR with positive allosteric modulators may have deleterious effects on the entire digestive system, possibly connected to the promotion of inflammatory processes and, by extent, of colitis associated colorectal cancer. More research will be necessary to define the mechanisms that underlie the effects of pharmacological calcimimetics and the CaSR on gastrointestinal inflammation and, in general, gastrointestinal wellbeing.

## 5. Conclusions

CaSR stimulation has profound effects on differentiation and inflammation in colon cancer cells overexpressing the CaSR. While the down-regulation of certain differentiation markers seemed to be cell line-specific, CaSR stimulation promoted the expression of pro-inflammatory markers in both studied cell lines, consistent with studies of a colitis model, and reports on lung inflammation, and warnings connecting CaSR stimulation with (gastrointestinal) inflammation. Indeed, these findings emphasize a need for caution in the clinical use of type II calcimimetics, especially for patients who suffer from inflammatory bowel diseases. Whether pharmacological inhibition of the CaSR could provide a new therapeutic approach for the treatment of IBDs requires further research.

## Supplementary data

Supplementary information is listed in the **Supplementary figures file: Figure S1**: Graphs representing the FACS sorting of the transduced cells. **Figure S2**: CaSR mRNA and protein expression in transduced Caco2 cells. **Figure S3**: Functional analysis of the transduced HT29 cells. **Figure S4**: Expression of inflammatory markers in transduced Caco2 cells. **Figure S5**: Expression of intestinal markers in transduced Caco2 cells. **Figure S6**: BrdU and MTT assay on transduced HT29 and Caco2 cells treated with allosteric CaSR modulators.

Supplementary information regarding the experiments performed in this work is listed in the **Supplementary materials and methods file**: **Intracellular calcium measurements:** description of the procedure to measure intracellular calcium release upon CaSR stimulation. **IP-one accumulation assay:** protocol describing an indirect estimation of IP3 synthesis by measuring IP1 level. **MTT assay:** protocol describing how we measured cell viability. **BrdU assay:** protocol describing how we measured cell proliferation. The sequences of primers used for the SYBR-green RT-qPCR reaction are listed in the **Supplementary Table S1**.

## List of abbreviations

GPCR: G protein-coupled receptor
DSS: dextran sulfate sodium
AOM: azoxymethane
FDA: Food and Drug Administration
MTT: 3-(4,5-dimethylthiazol-2-yl)-2,5-diphenyltetrazolium bromide
BrdU: bromodeoxyuridine
DAPI: 14’,6-diamidino-2-phenylindole
FDR: false discovery rate
GO: Gene Ontology
G_q/11_: G protein subunit alpha q/11
PLC: phospholipase C
IP3: inositol 1,4,5-trisphosphate
IP1: inositol monophosphate
IL-1α: interleukin-1 alpha
IL-6: interleukin-6
IL-8: interleukin-8
IL-23α: interleukin-23 alpha subunit
CCL-20: C-C motif chemokine ligand 20
CSF1: colony stimulating factor 1
IL-34: interleukin-34
COX2: cyclooxygenasese 2
PDL-1: programmed cell death 1
MYO7B: myosin 7 b
ANKS4B: Ankyrin repeat and sterile alpha motif domain containing 4B
USH1C: Usher syndrome 1 c
CDX2: caudal type homeobox 2
NEUROG3: neurogenin3
ATOH1: atonal BHLH transcription factor 1
DLL1: delta like canonical NOTCH ligand 1
DLL4: delta like canonical NOTCH ligand 4
PECAM1 or CD31: platelet and endothelial cell adhesion molecule 1
NF-κB: nuclear factor kappa B
IBD: inflammatory bowel diseases
INSM1: insulinoma-associated 1
PAX4: paired box 4
MUC2: mucin 2

## Availability of data and materials

The dataset generated by the RNA-seq in the current study is available in the GEO repository: GSE140984, accessed on 05/12/2019 at the following link https://www.ncbi.nlm.nih.gov/geo/query/acc.cgi?acc=GSE140984

## Author’s contribution

Conception and design of the project: L.I. and E.K.; execution of the experiments and data collection and interpretation: L.I., T.E., K.G., J.D., T.M., M.S., S.D. and E.K.; animal experiments: L.I., P.H., T.E. and M.S.; manuscript writing/editing L.I., T.E., P.H., M.G., S.D., S.B.P., M.S. and E.K. All authors approved the final version of the current manuscript.

## Acknowledgements

We would like to express our gratitude to the following people: to Martina Salzman that performed the paraffin-embedding and microdissection of the tumor biopsies; to Mag. Gerhard Zeitler, the technician of the animal facility that took care of the mice executing also the gavages and the tumor measurements; to Mag. Gerald Timelthaler and Dr. Karin Schelch who provided theoretical and technical support for the use of the confocal microscope. We would like to thank The Genomics Core Facilities of the Medical University of Vienna, a member of VLSI, for sequencing and initial data analysis of the RNA-seq and to thank Professor Andreas Spittler and Ing. Günther Hofbauer from the Flow Cytometry Core Facilities of the Medical University of Vienna for performing the FACS sorting. Finally, we would like to thank Professor Arthur Conigrave for critically reviewing the manuscript.

