## supplementary figures for "Effects of pharmacological calcimimetics on colorectal cancer cells over-expressing the human calcium-sensing receptor"

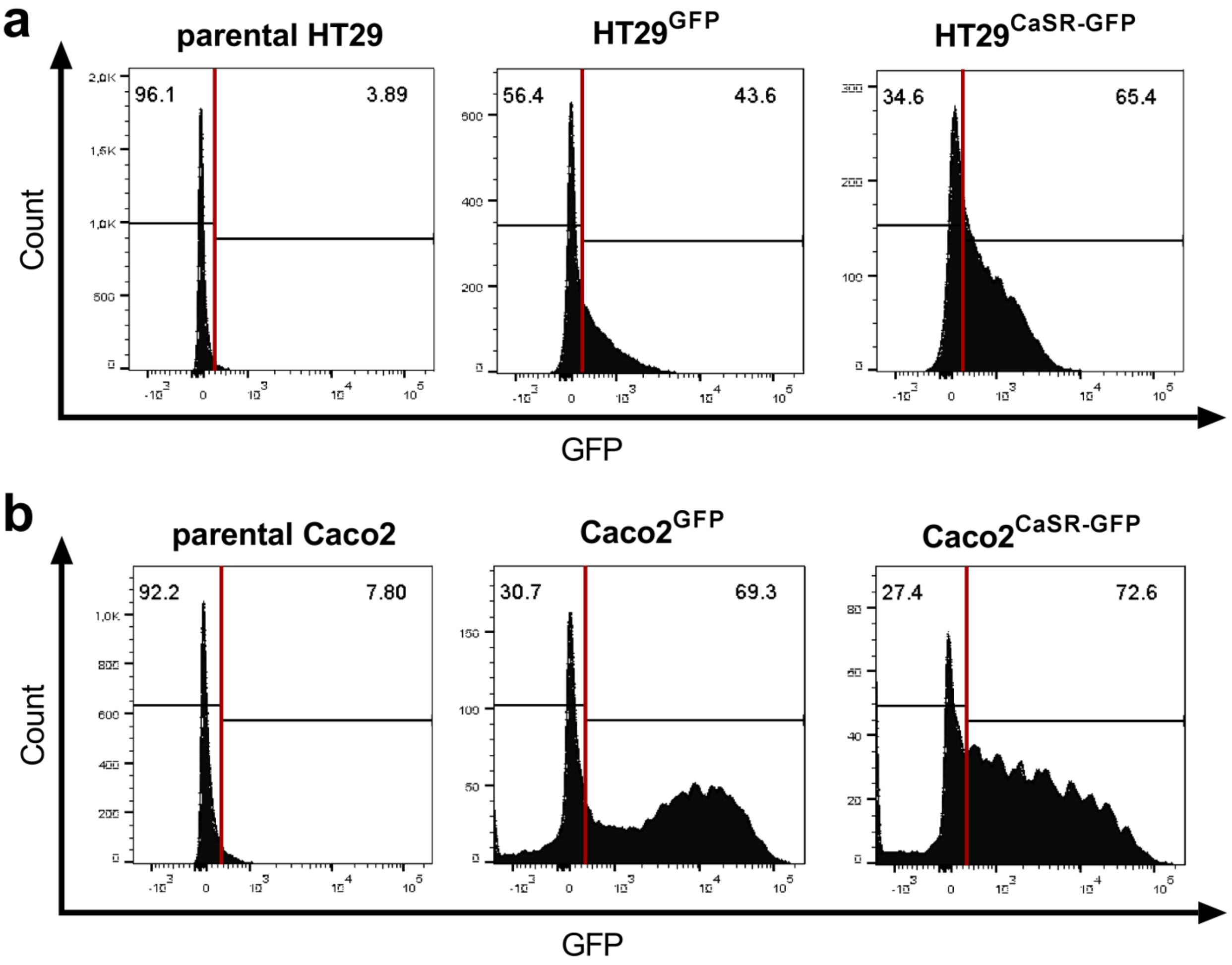


**Fig. S1 FACS sorting of the transduced cells.**
The transduced HT29 (**a**) and Caco2 (**b**) cells were sorted based on GFP intensity, the parental cell lines (not transduced), were used to measure the fluorescence background. The numbers on the top-right of the graphs represent the percentages of GFP-positive cells relative to the total cell population, while the numbers on the top-left represent the percentages of GFP-negative cells.

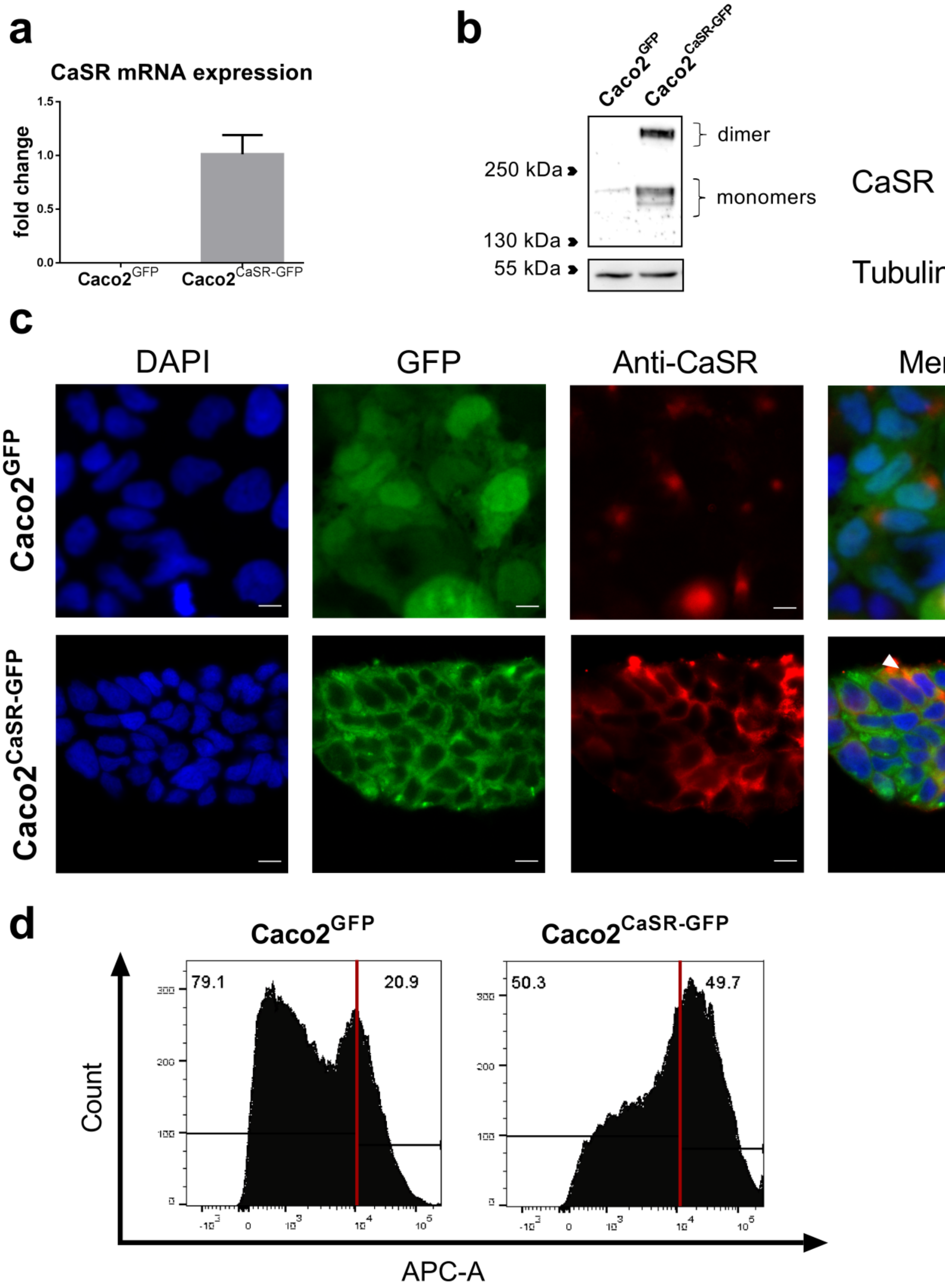


**Fig. S2 CaSR expression in transduced Caco2 cells.**
CaSR mRNA quantification, values normalized to the mean of CaSR expression in Caco2$^{CaSR\text{-}GFP}$ cells + SD (n=3) (**a**). Representative western blot for CaSR protein expression; in Caco2$^{CaSR\text{-}GFP}$ cells the CaSR monomers and the dimer corresponding to the CaSR-GFP fusion-protein were detected (**b**). Representative confocal images of GFP (green) and CaSR (red) (immuno-)fluorescence; white arrows indicate the points of co-localization (yellow/orange); scale bars represent 10µm (**c**). Representative histograms showing the percentages of CaSR-negative (left) and –positive (right) cells detected via flow cytometry, using the allophycocyanin (APC-A) channel (**d**).

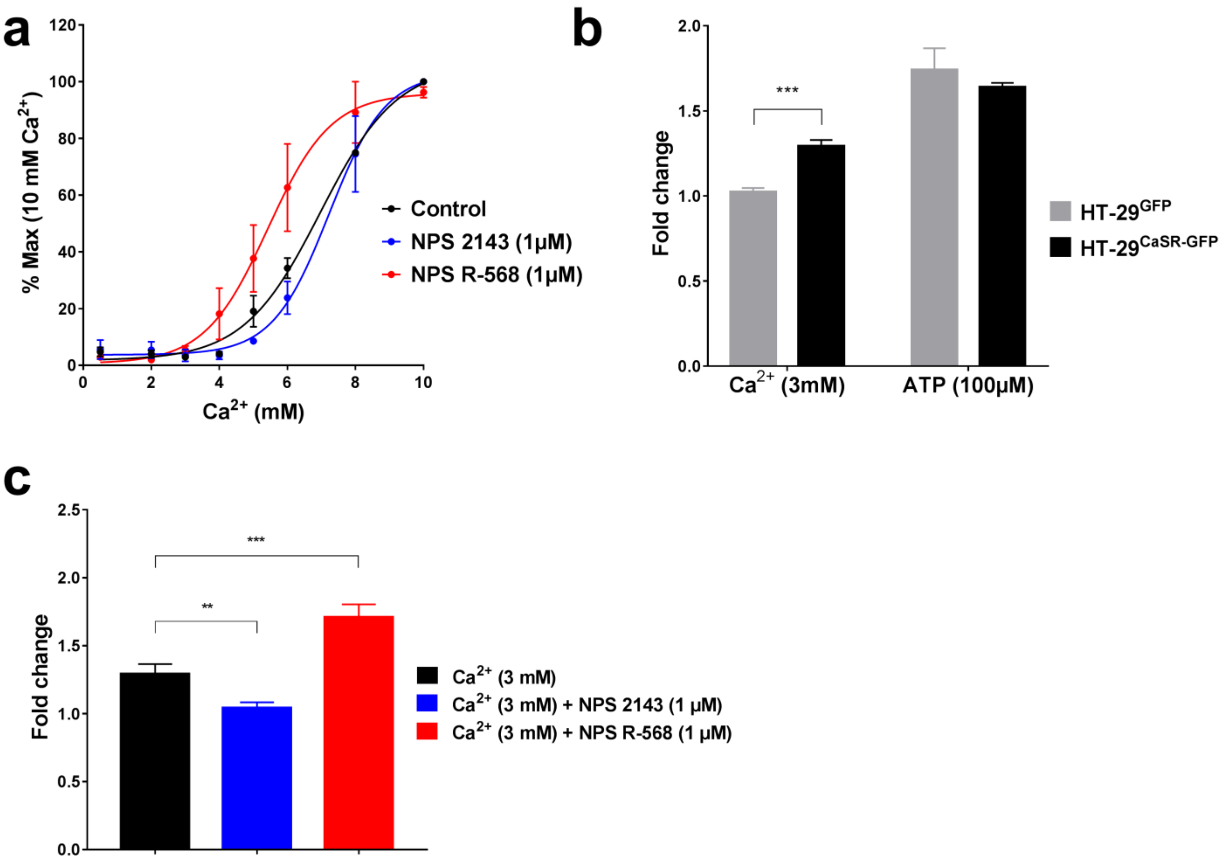


**Fig. S3 CaSR functionality of HT29 transduced cells.**
IP1 secretion in HT29$^{CaSR\text{-}GFP}$ cells treated with 0.1% DMSO (vehicle control) or with allosteric CaSR modulators under increasing concentrations of extracellular calcium (**a**). Intracellular $Ca^{2+}$ mobilization in HT29$^{CaSR\text{-}GFP}$ and HT29$^{GFP}$ cells was measured after stimulation with 3mM extracellular $Ca^{2+}$; the values are expressed as fold change relative to the baseline (pre-incubation with 0.5mM $Ca^{2+}$). 100µM ATP was used as technical positive control (**b**). Intracellular $Ca^{2+}$ mobilization in HT29$^{CaSR\text{-}GFP}$ cells in the presence of vehicle or allosteric CaSR modulators with 3mM extracellular $Ca^{2+}$ (**c**). Data are presented as mean ± SD (n=3). Statistical analysis was performed using unpaired t-test (**b**) and one-way ANOVA with Dunnett's post-hoc test (**c**), **** $p<0.0001$; *** $p<0.001$; ** $p<0.01$; * $p<0.05$.

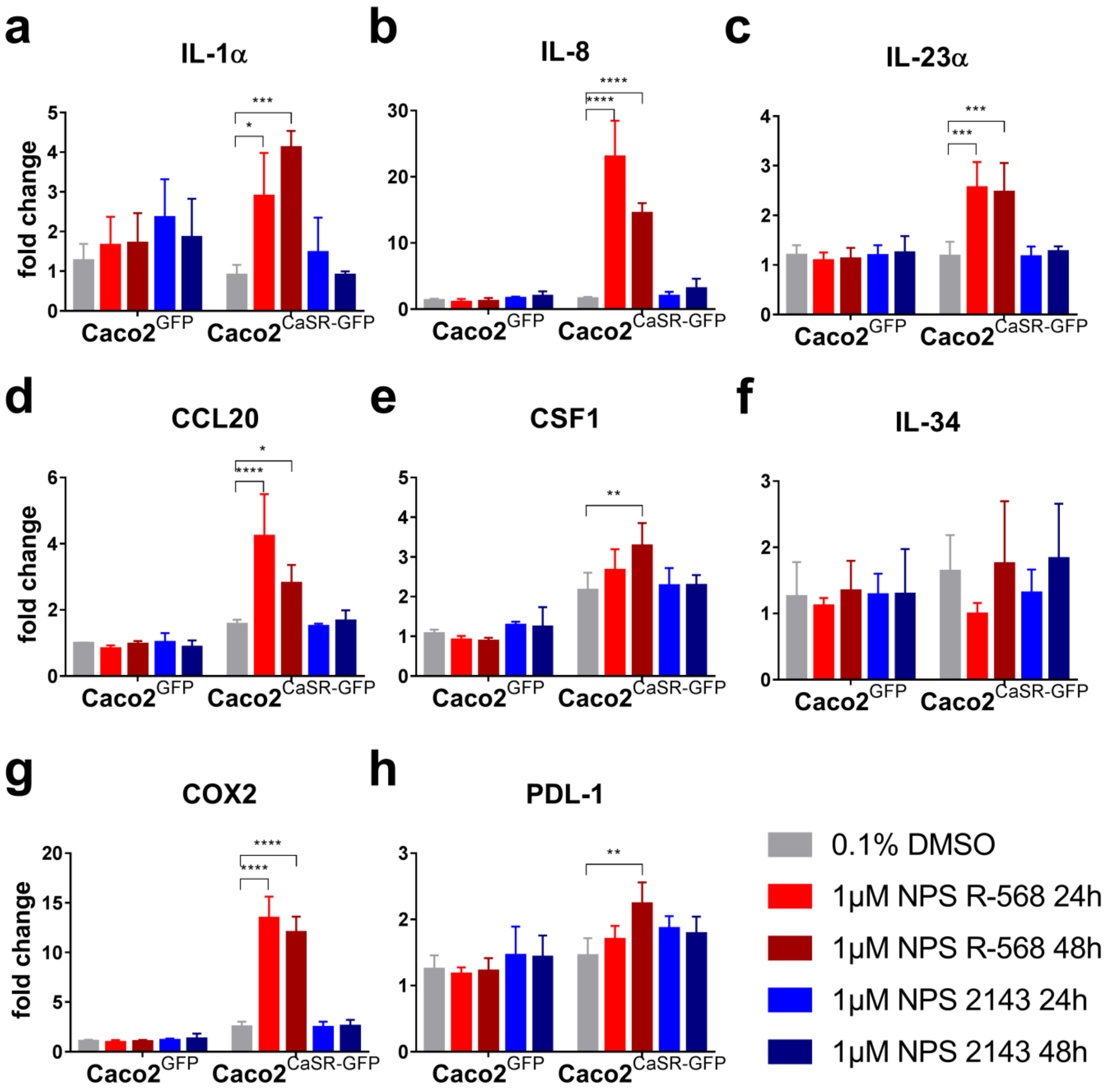


**Fig. S4 Expression of inflammatory markers in transduced Caco2 cells.**
RT-qPCR analysis of the mRNA level of IL-1α (**a**), IL-8 (**b**), IL-23α (**c**), CCL20 (**d**), CSF1 (**e**), IL-34 (**f**), COX2 (**g**) and PDL-1 (**h**). All values were normalized to the untreated Caco2$^{GFP}$ cell as control. Data are presented as mean + SD (n=3). Statistical analysis was performed using two-way ANOVA with Dunnett's post-hoc test, **** p<0.0001; *** p<0.001; ** p<0.01; * p<0.05.

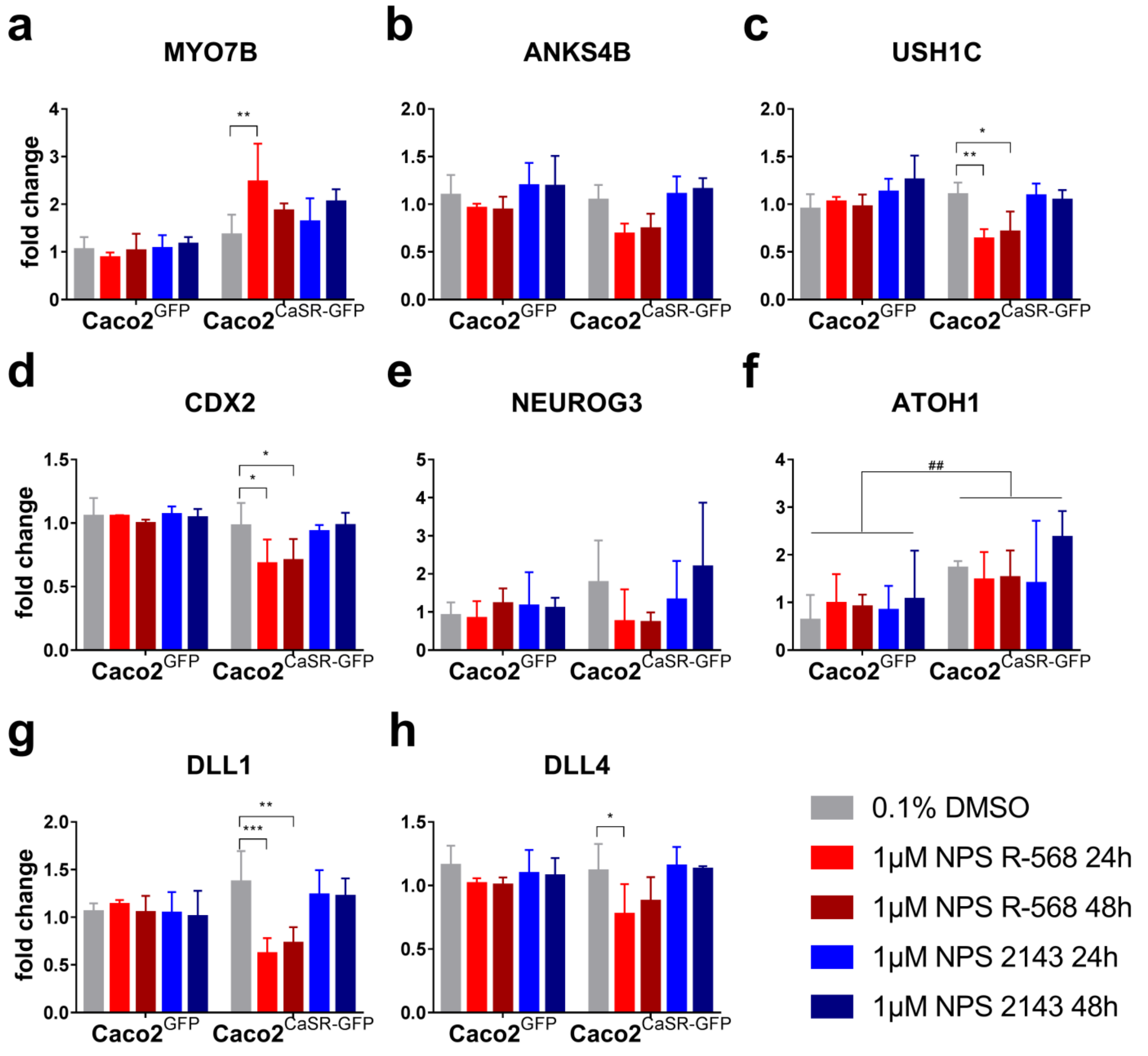


**Fig. S5 Expression of intestinal markers in transduced Caco2.**
RT-qPCR analysis of the mRNA level of MYO7B (**a**), ANKS4B (**b**), USH1C (**c**), CDX2 (**d**), NEUROG3 (**e**), ATOH1 (**f**), DLL1 (**g**) and DLL4 (**h**). All values were normalized to the untreated Caco2$^{GFP}$ cell as control. Data are presented as mean + SD (n=3). Statistical analysis was performed using two-way ANOVA with Dunnett's post-hoc test, **** $p<0.0001$; *** $p<0.001$; ** $p<0.01$; * $p<0.05$; $^{####}$ $p<0.0001$; $^{###}$ $p<0.001$; $^{##}$ $p<0.01$; $^{\#}$ $p<0.05$.

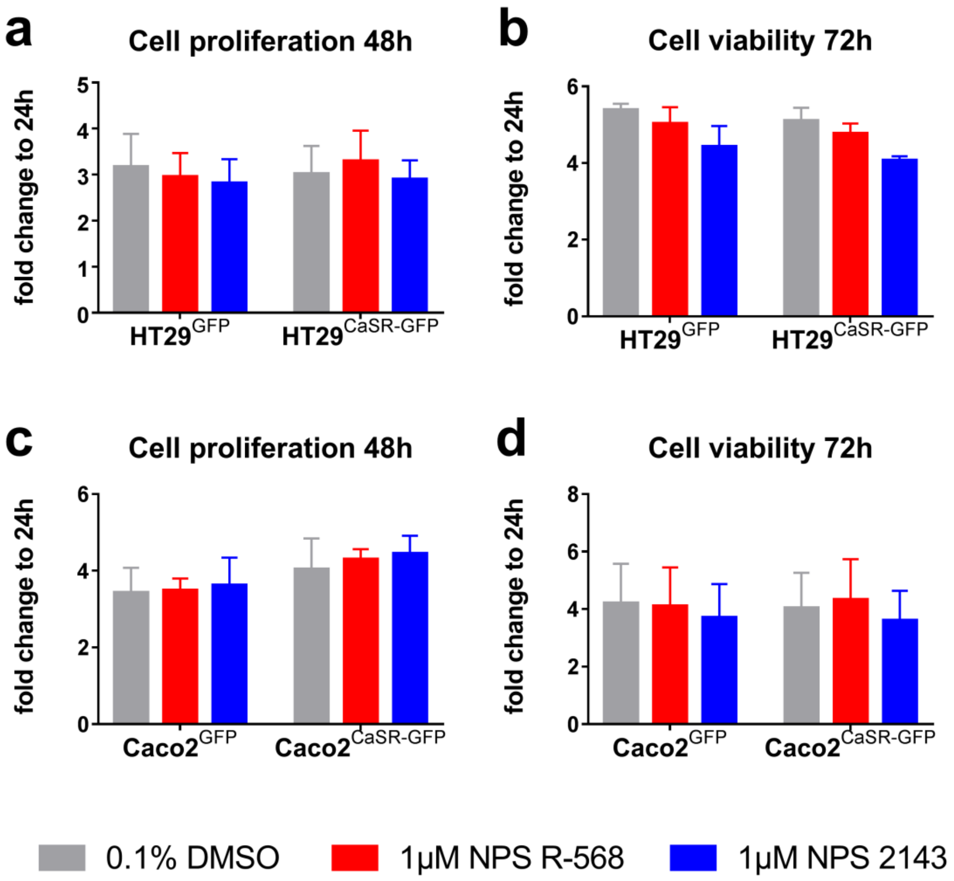


**Fig. S6 Cell proliferation and viability.**
Cell proliferation (BrdU incorporation assay) of transduced HT29 and Caco2 cells treated for 48h with CaSR-modulators (**a, c**). Cell viability (MTT assay) of transduced HT29 and Caco2 cells treated for 72h with CaSR-modulators (**b, d**). Data are presented as mean + SD (n=3) of fold change relative to 24h post-seeding plate (normalizing plate) to minimize seeding errors. Statistical analysis was performed using two-way ANOVA with Dunnett's post-hoc test.
