## supplementary materials and methods for "Effects of pharmacological calcimimetics on colorectal cancer cells over-expressing the human calcium-sensing receptor"

### Intracellular calcium measurements

Stably transduced HT29 cells were seeded onto 13 mm glass coverslips and cultured until fully confluent. On the day of the experiment, each coverslip was washed with assay buffer ($Ca^{2+}$ and $Mg^{2+}$ free HBSS, 20mM HEPES, 0.5mM $Ca^{2+}$) and then they were incubated in assay buffer containing 2μM FURA-2-AM pre-mixed with pluronic F127 (Sigma-Aldrich), and 33mM glucose, for 2.5 hours at room temperature. The coverslips were then placed into a glass cuvette filled with assay buffer and the fluorescence was measured with an F-4500 fluorescence spectrophotometer (Hitachi, Japan). CaSR modulators were added directly into the cuvette and mixed for 1 minute before fluorescence recording. Emissions of calcium-bound and unbound FURA-2 were measured at 340 and 380 nm excitation respectively. Raw data were analyzed using FL-solutions 2.0 software (Hitachi).

### IP-one accumulation assay

The IP-One kit (CisBio, France) was used to measure inositol monophosphate (IP1) levels in transduced HT29 cells. $4 \times 10^4$ cells were seeded in 48-well plates and grown for 4 days. On the day of the experiment, cells were washed with assay buffer containing $Ca^{2+}$- and $Mg^{2+}$-free HBSS, 20mM HEPES adjusted to pH 7.4. After washing, the cells were stimulated for 30 minutes at 37 °C with 1μM of allosteric CaSR modulators (see results) mixed in ligand buffer ($Ca^{2+}$- and $Mg^{2+}$-free HBSS, 20mM Hepes, 20mM LiCl, adjusted to pH 7.4). This buffer was then discarded and the cells were washed once again with assay buffer and lysed with IP-one lysis and detection buffer (CisBio) for 30 minutes at room temperature. 14 μL of this cell lysate were then transferred into 96-well low volume plates (CisBio), and 6 μL of pre-mixed detection solution (anti-IP1-cryptate Terbium conjugate and IP1-d2 conjugate) were added to all wells containing sample and standards. The plate was then sealed and incubated for 1 hour at room temperature in the dark. Samples were excited with light at 340nm and fluorescence was measured at 665nm and 620nm wavelength using the Infinite M1000 Pro (Tecan, Austria). The homologous time-resolved fluorescence (HTRF) ratio (665/620), which is inversely proportional to IP1 accumulation, was used to determine the IP1 response. HTRF ratios were extrapolated to IP1 concentrations from standard curves obtained from the IP1 standards (CisBio).

**MTT assay**

$5x10^3$ cells were seeded in 96-well plates and cultured within the first 24 hours in normal medium, 1 plate for normalization (24h post seeding time point) and 1 plate for the treatments (experimental plate). After 24h, we started the treatments on the experimental plate. To perform the MTT assay the cells were incubated in DMEM + 10% FCS + 1mg/ml thiazolyl blue tetrazolium bromide (Sigma-Aldrich) for 2h in the incubator. The reaction produced a reduced formazan salt that was solubilized in 50% isopropanol + 50% DMSO solution. Absorbance was measured at 570nm with Infinite M200 Pro (TECAN) microplate reader.

**BrdU assay**

$1x10^3$ cells were seeded in black 96-well pates with transparent flat bottom and cultured within the first 24 hours in normal medium, 1 plate for normalization (24h post seeding time point) and 1 plate for the treatments (experimental plate). After 24h, we started the treatments on the experimental plate. Cell proliferation was assessed using “Cell Proliferation ELISA, BrdU chemiluminescent” assay (Roche, Switzerland) following the manufacturer’s protocol. Chemiluminescence was recorded with Infinite M200 Pro (TECAN) microplate reader.
