## supplementary table 1 for "Effects of pharmacological calcimimetics on colorectal cancer cells over-expressing the human calcium-sensing receptor"

**RT-qPCR primers list**

| Primer target | forward | reverse |
|---|---|---|
| CSF1 | GGAGACCTCGTGCCAAATTA | TATCTCTGAAGCGCATGGTG |
| USH1C | CGGCTCCTACGCATCAAGAA | GCCAGGGTGTAGTCTGTCAC |
| ANKS4B | AAGAGGCTACCAAGCGAGAT | TTCCCCAGATGTCACACCTA |
| MYO7B | TGAGACGCACCTATGCCAAT | GGATGGGGATGTGCTTCTTG |
| IL-34 | TTGACGCAGAATGAGGAGTG | CCCTCGTAAGGCACACTGAT |
| IL-23A | CTCAGGGACAACAGTCAGTTC | ACAGGGCTATCAGGGAGCA |
| CCL20 | TTGCTCCTGGCTGCTTTGAT | GCCGTGTGAAGCCCACAATA |
| IL-1A | GGTTGAGTTTAAGCCAATCCA | TGCTGACCTAGGCTTGATGA |
| CD274 (PDL-1) | TATGGTGGTGCCGACTACAA | TGCTTGTCCAGATGACTTCG |
| PTGS2 (COX2) | CAAGACAGATCATAAGCGAGGG | GTCTAGCCAGAGTTTCACCG |
| DLL1 | ACCTGCGAGCTGGGGATTGA | AGCGGCACAGGTAGGCATCA |
| DLL4 | AACTGCCCTTCAATTTCACCT | GCTGGTTTGCTCATCCAATAA |
| ATOH1 | CAGCTGCGCAATGTTATCCC | TTGTAGCAGCTCGGACAAGG |
| NEUROG3 | AGCCGGCCTAAGAGCGAGTT | TTGGTGAGCTTCGCGTCGTC |
| IL-8 | CTTGGCAGCCTTCCTGATTT | TTCTTTAGCACTCCTTGGCAAAA |
| CDX2 | AGGGGGTGGTTATTGGACTC | CATTCAGCCCAGAGAAGCTC |
| RPLP0 | CCTCATATCCGGGGGAATGTG | GCAGCAGCTGGCACCTTATTG |

**TableS1.** List of forward and reverse sequences of the primers for the SYBR-green RT-qPCR reaction.
